# A Boltzmann model predicts glycan structures from lectin binding

**DOI:** 10.1101/2023.06.03.543532

**Authors:** Aria Yom, Austin Chiang, Nathan E. Lewis

## Abstract

Glycans are complex oligosaccharides involved in many diseases and biological processes. Unfortunately, current methods for determining glycan composition and structure (glycan sequencing) are laborious and require a high level of expertise. Here, we assess the feasibility of sequencing glycans based on their lectin binding fingerprints. By training a Boltzmann model on lectin binding data, we predict the approximate structures of 88 *±* 7% of N-glycans and 87 *±* 13% of O-glycans in our test set. We show that our model generalizes well to the pharmaceutically relevant case of Chinese Hamster Ovary (CHO) cell glycans. We also analyze the motif specificity of a wide array of lectins and identify the most and least predictive lectins and glycan features. These results could help streamline glycoprotein research and be of use to anyone using lectins for glycobiology.

## Introduction

Glycans are diverse oligosaccharides found in every branch of life. In humans, they are critical to protein folding, immunity, cell-cell interactions, and many other processes. Improper glycosylation is implicated in several disease states, and may be useful as an early indicator of cancer.^1–6^ Proper glycosylation is vital to the efficacy of many protein therapeutics and vaccines.^7–9^ The influence of glycans on protein function and organism physiology highlights the importance of finding flexible and affordable methods for glycan sequencing.

Currently, the most common method used to determine the structure of a glycan is to use mass spectrometry (MS) coupled with expert annotation, or potentially automated annotation.^10,11^ Other experimental techniques also exist, such as NMR^12,13^ and exoglycosidase treatment.^14,15^ Unfortunately, these methods are expensive and require a high level of expertise to conduct the experiments and to interpret the results. Also, with mass spectrometry, it can be difficult to distinguish bond orientations, which can impact physiological responses.^16^ Lectins, antibodies, and other carbohydrate-binding proteins display stereospecific binding, and many glycan-binding proteins (GBPs) can be readily obtained from commercial sources.^17^ Thus, they have long been employed to quickly discern the presence or absence of certain glycan motifs.^18^ Arrays of lectins have been used to generate binding profiles for glycoproteins, and to assay the corresponding cellular interactions.^19–22^ GBPs have also been used to augment MS data for automated glycan sequencing;^23^ however, the challenge of sequencing glycans from GBPs alone has not been addressed.

Recent studies have utilized machine learning (ML) techniques to extract the binding specificities of a plethora of commonly used lectins,^24–27^ demonstrating the utility of ML for lectin analytics. ML has also been used for other tasks in glycobiology, such as glycan classification and glycosylation site prediction.^28–32^ We refer the reader to references [^33–35^] for more comprehensive reviews of the present and future of ML and big data in glycobiology. Here we approach the problem of *de novo* glycan sequencing, i.e., the elucidation of the composition and structure an unknown glycan. Though a single lectin clearly cannot extract as much information as an LC-MS experiment, it is not clear how much information can be obtained from an array of lectins. One may therefore ask: by leveraging machine learning, can one determine the structure of a glycan from its lectin binding fingerprint alone?

In this study, we show that a simple Boltzmann model using a wide panel of lectins can determine the approximate structures of 88 *±* 7% of N-glycans and 87 *±* 13% of O-glycans from the Consortium for Functional Glycomics (CFG) glycan arrays. We also demonstrate that our model can sequence glycans from Chinese Hamster Ovary (CHO) cells, the cell type of choice for producing most therapeutic glycoproteins.^36^ Then, using information theoretic measures, we identify the most robust lectin-motif binding pairs, which we list as a resource for those using lectins in their research. We also determine which glycan motifs are most elusive to currently available lectins, thus suggesting profitable future directions for the engineering of new GBPs.

## Results

We first obtained and analyzed lectin binding data from CFG glycan arrays^24^ to test if a machine learning model can accurately extract and encode the motif-binding patterns of each lectin. We then validated the accuracy of our model on both the CFG glycan set and a comprehensive set of CHO N-linked glycans from two previous studies.^24,37^ The results of these analyses follow.

### A Boltzmann model is near-optimal for encoding lectin binding

In what follows, we model each glycan as a tree with monosaccharides as nodes and bonds as edges. We then define a motif as being any connected subgraph of such a glycan, such as a fucosylated N-core (see Methods - Motif recognition for more details). Consistent with previous studies, we find that each lectin has a small set of motifs to which it binds.

Since lectins are not expected to have logically complex binding rules, we expect shallow neural network topologies to be adequate. A reasonable minimal model for this interaction would be a fully visible Boltzmann machine, which can be conceptualized as a two layer neural network (Figure 1). In this model, we generate a likelihood for each possible glycan, based on the lectin binding signal received.

**Figure 1:**
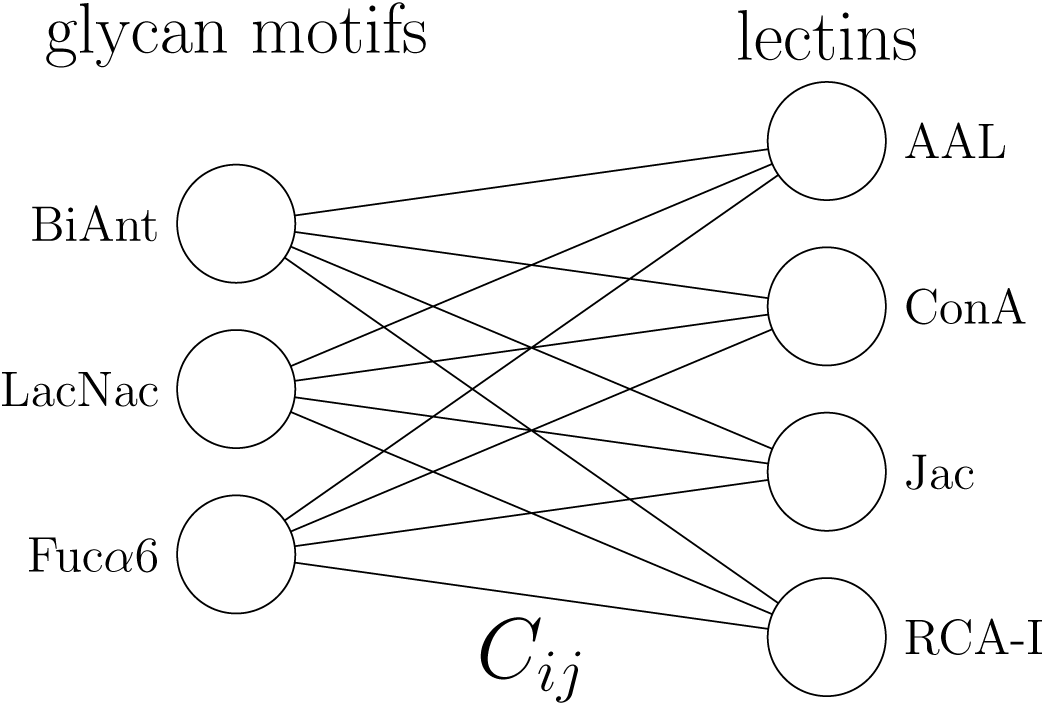
Boltzmann model

Boltzmann Model:

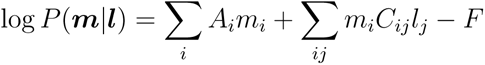

where {*m_i_*} and {*l_i_*} are binaries representing the presence of a motif or the binding of a lectin, respectively, *A* and *C* are the model parameters, and *F* is a normalization constant.

Could a more complex machine learning architecture (e.g., a deep neural network) outperform such a simple model? This would be true if, for example, a lectin binds a glycan only when two motifs are present, but not when either one is present, causing the likelihood function to be nonlinear. The key property of such a scenario is the presence of higher order interactions, i.e., those involving the product of more than one lectin or more than one motif. One way to parameterize a maximally general model would be to include all higher order interactions. The next level up from our Boltzmann model would be to include only the three-party interactions, i.e. those between two lectins and one motif, and those between two motifs and one lectin.

Second-Order Model:

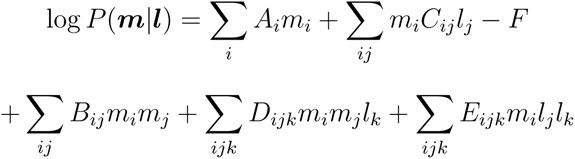

where {*m_i_*} and {*l_i_*} are binary variables representing the presence of a motif or the binding of a lectin, respectively, *A*-*E* are the model parameters, and *F* is a normalization constant.

One way to gauge the utility of adding these extra terms would be to measure the impact of these terms on the amount of information extracted about any one motif. This can be quantified using the cross entropy Δ*H* (see Methods - Cross entropy). In general, we find that our minimal model is virtually indistinguishable from the Second-Order model (Figure 2). Therefore, if we assume similar diminishing gains from adding further higher order terms, we can conclude that our simple Boltzmann model is close to optimal at encoding lectin binding patterns.

**Figure 2:**
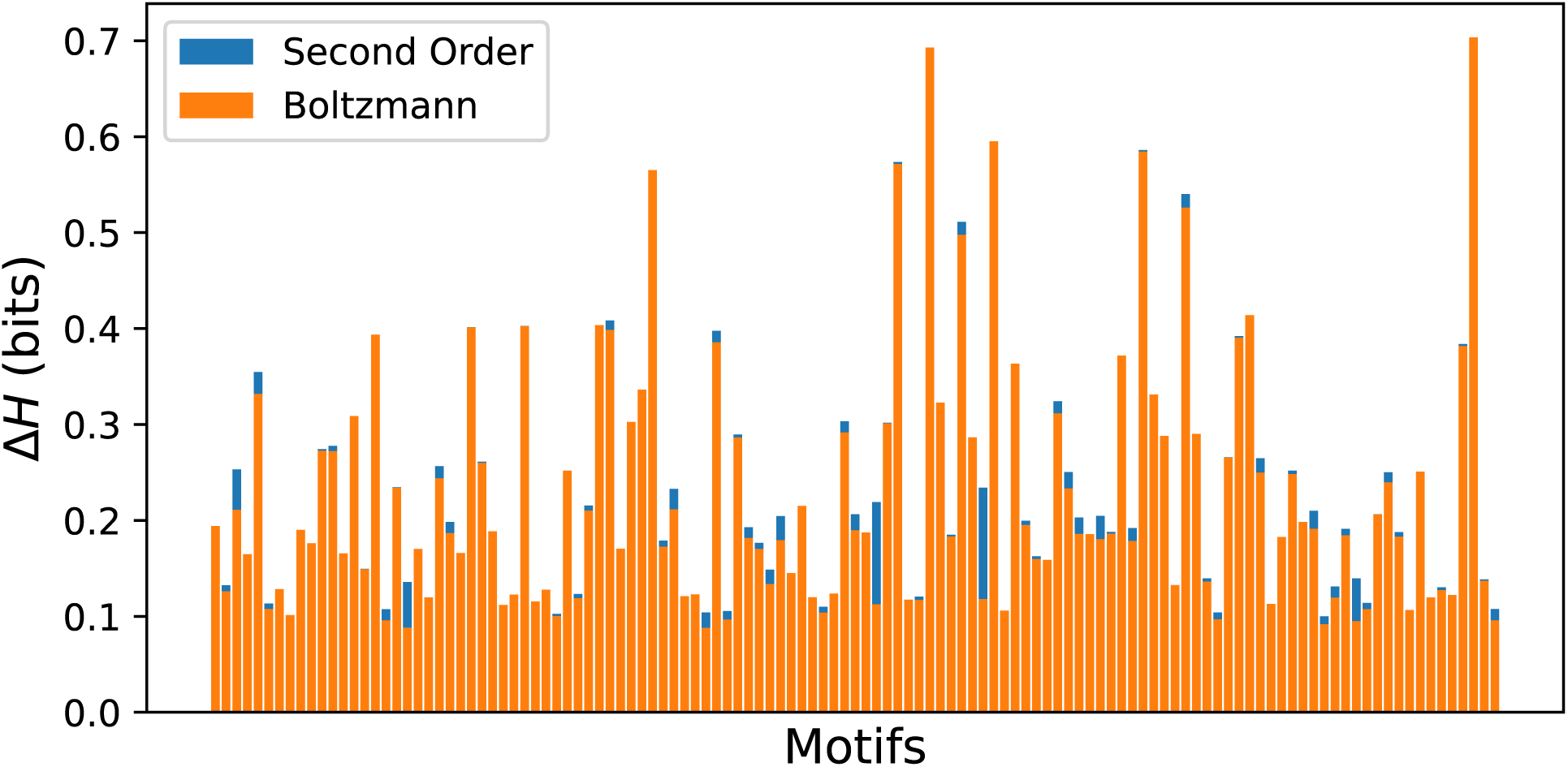
Amount of information extracted about each motif by our first and second order models, as quantified by the difference in cross entropy Δ*H*. For each motif, the models are optimized over the two most predictive lectins. For the task of predicting any single motif, there is little to be gained from using the higher order model. See Methods - Cross entropy for more details.

**Figure 3:**
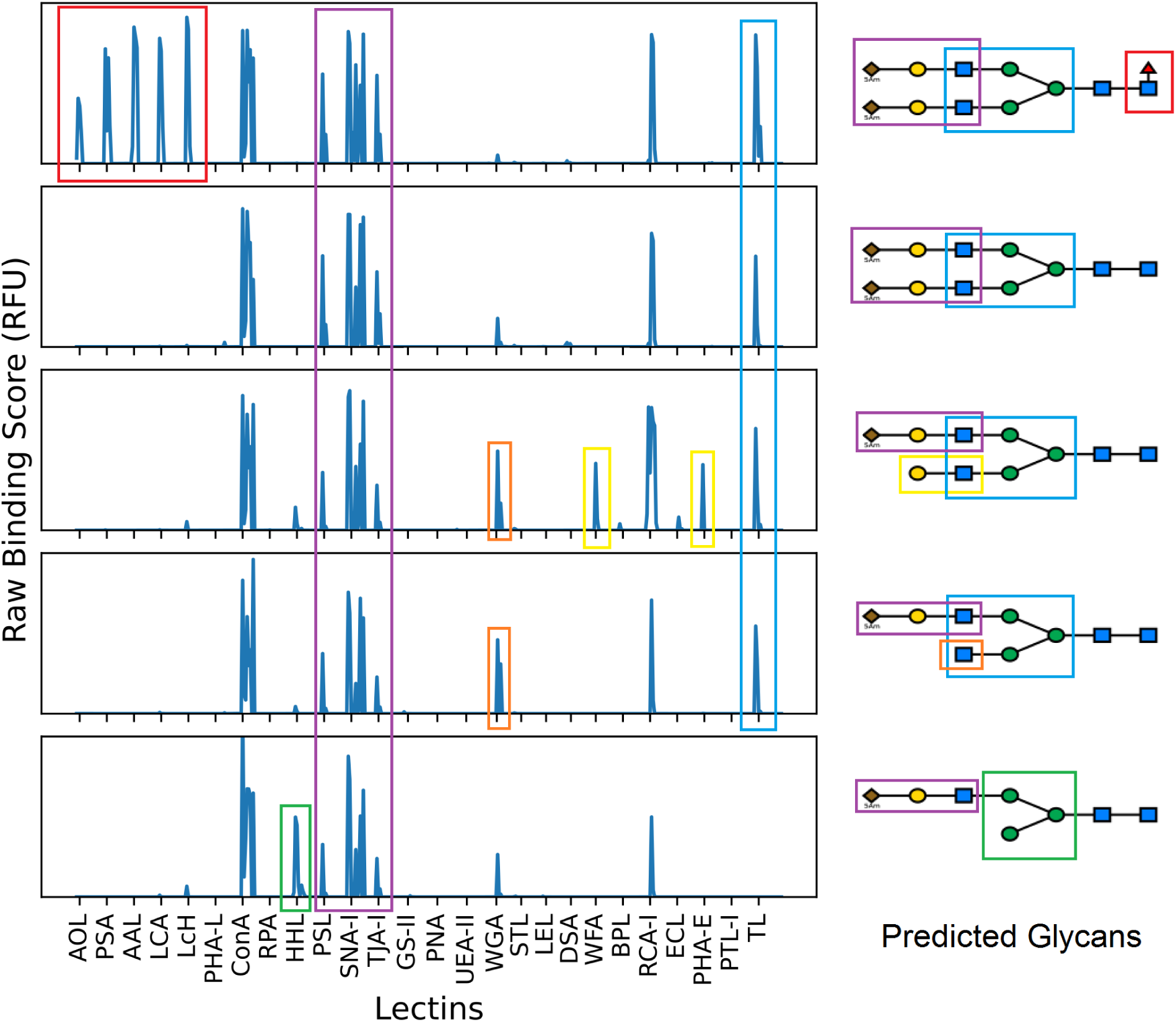
Conceptual illustration of how prediction occurs on five slightly different glycans. The model associates each lectin with certain features. The model then finds glycans that possess these features. The model also learns to ignore certain features at times, like the WGA peak in the center glycan. For each lectin, several concentrations were used, producing different degrees of binding, though only the lectin code is labelled along the x-axis.

### The model predicts glycan structures from lectin profiles

Given binding measurements from a panel of lectins, our model will predict the structures of glycans being bound. Since our model is probabilistic, each glycan will be predicted with an associated likelihood. These likelihoods can then be used to rank the glycans, with rank 1 glycans being the model’s best prediction, rank 2 being the second most likely glycan, and so on. We are interested not only in how often our model predicts the exact correct glycan, but also in how often the correct glycan is among its top *n* predictions.

We begin by generating a set of glycans sufficiently large and diverse as to encompass all CFG glycans and all other glycans that could plausibly exist given the linkage patterns found in the CFG dataset. Our approach is analogous to the “virtual glycome database” of [^23^], but in the interest of *de novo* sequencing, we do not manually input the set of possible patterns. Instead, our algorithm infers a minimal Markov model from the input glycans and then samples it to generate a broad range of plausible glycans (see Methods – Markov modelling of glycan space). We find that the model entropy is 12 bits, so only need on the order of 10^4^ glycans to cover the full set of plausible glycans. For each lectin profile, a likelihood is calculated for all generated glycans, in addition to all CFG glycans. Since the predicted structures can come from a diverse set of glycans missing from the CFG array, the prediction can be regarded as an act of *de novo* glycan sequencing.

To test the model’s glycan prediction accuracy, we randomly separated the Consortium for Functional Glycomics (CFG) glycans into training and test sets using an 80/20 split, and then trained and tested our model on the corresponding binding data from 64 lectins and 9 antibodies.^24^ Our model was highly predictive on N-glycans, predicting the exact structure for 61 *±* 10% of glycans, and including the correct structure in its top three predictions for 88 *±* 7% of glycans (Figures 4, 7, and S-2).

**Figure 4:**
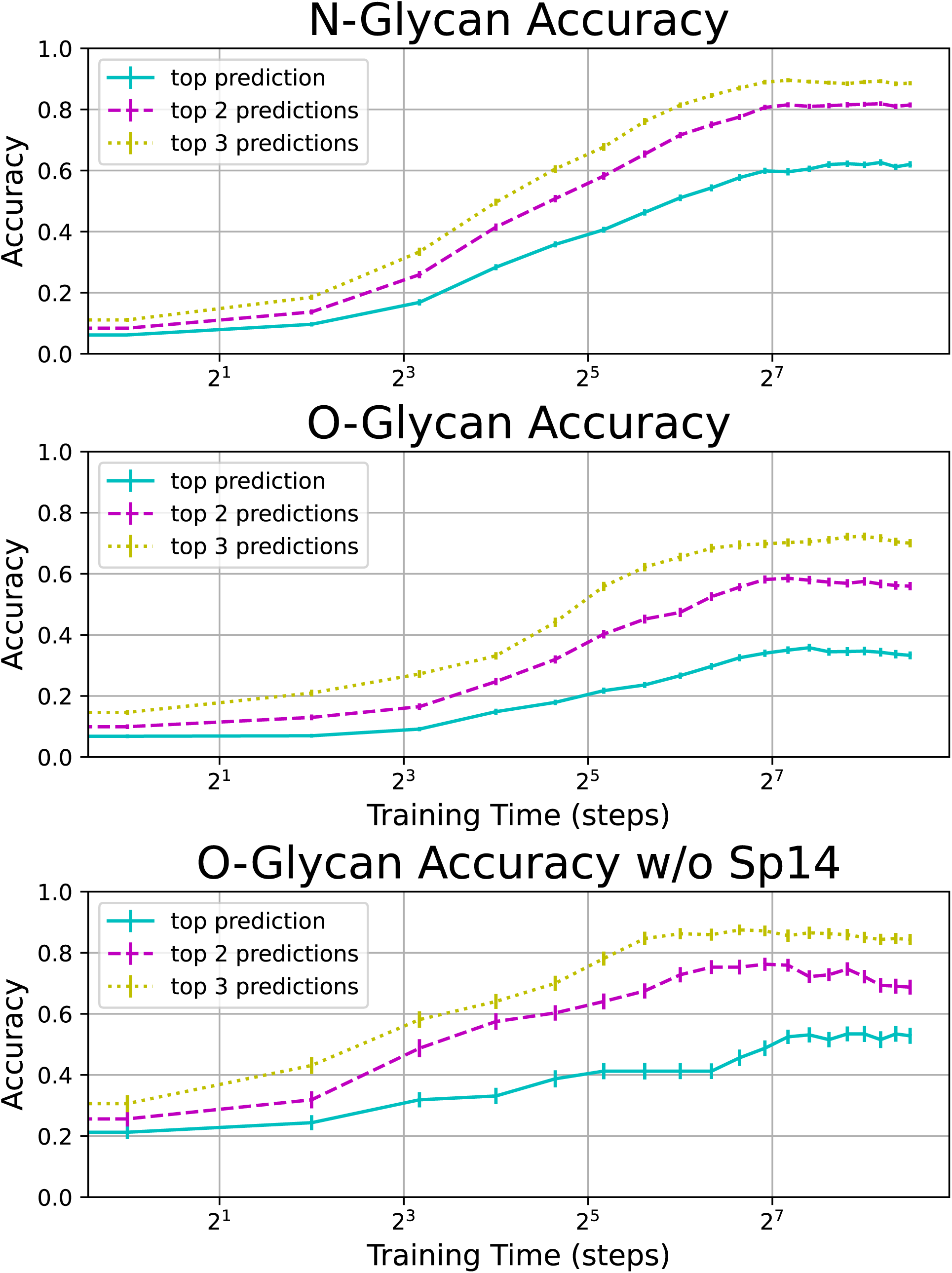
Test accuracy on N-linked and O-linked CFG glycans. Results averaged over 64 training/test set partitions. Error bars represent the standard error. Note that when the problematic spacer Sp14 is removed, O-glyca_9_n accuracy is much higher. A random sample of correct and predicted glycan structures may be found in Figure S-2.

Our model was less predictive on O-glycans, and we speculated that this may be due to improper presentation on the array. Since spacer 14 (Sp14) on the CFG array previously was shown to have a tendency to collapse O-glycans, we cleansed our data of Sp14. This resulted in a dramatic increase in O-glycan prediction accuracy to similar levels as our N-glycans (Figures 4, 7, and S-2). In concordance with an earlier study, ^38^ we did not find any N-glycan spacers with unusually low accuracy. We direct the interested reader to Figure S-2 for a large random sample of CFG glycans and their corresponding model predictions.

So which glycans were well predicted, and which were not? To answer this, we extracted three groups of glycans based on their ranks. Our high accuracy group contained only exactly predicted glycans, while our medium and low accuracy groups contained glycans with ranks 3 or 4 and ranks *>* 6, respectively. We did not find the accuracy to be dependent on size or number of branches; however, we did find certain motifs to be enriched in each group. By comparing the total number of occurrences of each motif to the number of occurrences in each group, we could perform a binomial test and compute a p-value for each motif and group. The lowest p-value motifs for each group can be found in Figure 5. Many of the high accuracy motifs are also sharply bound by lectins (Figure 8).

**Figure 5:**
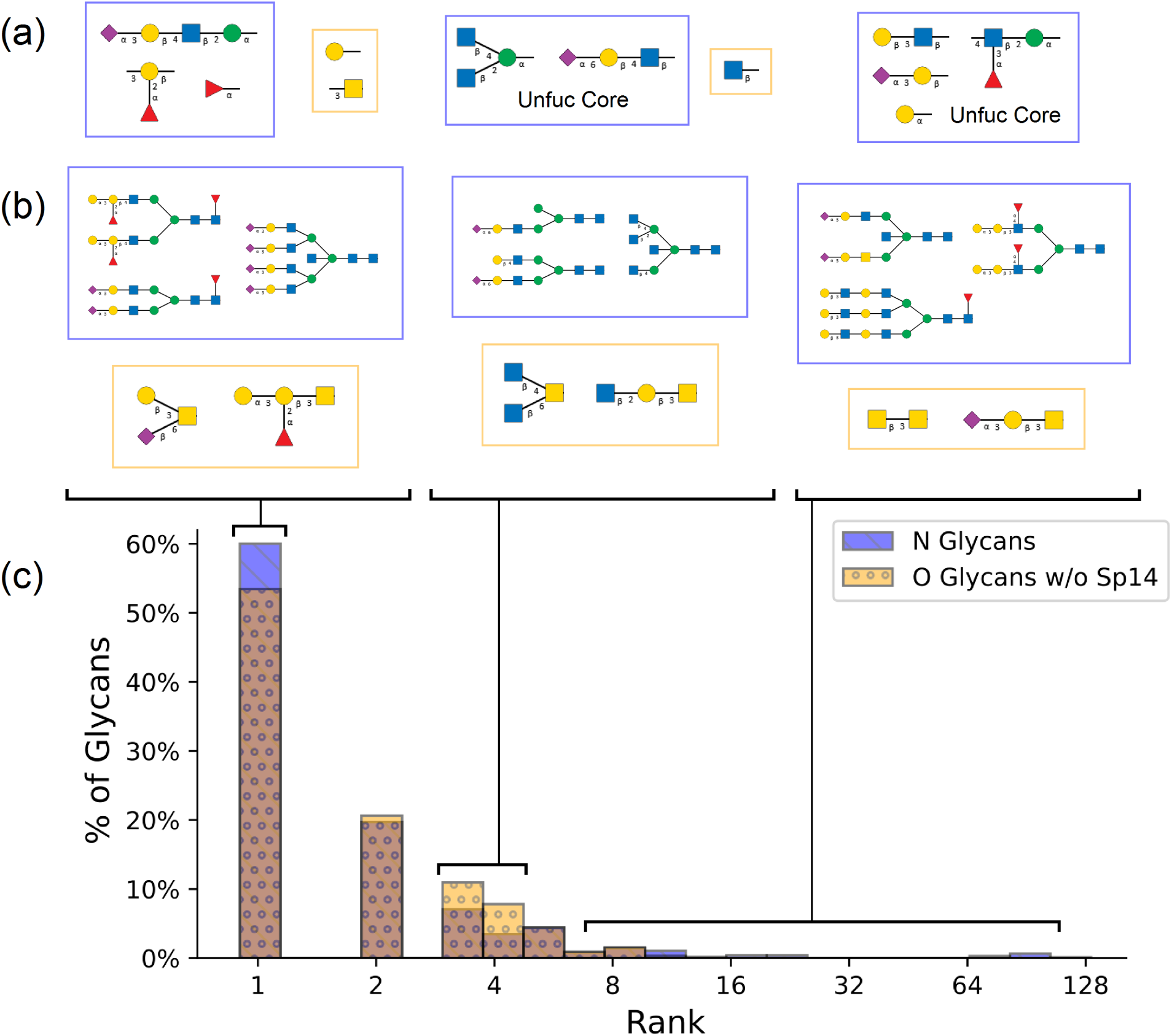
Using the prediction results from 64 runs of our algorithm with random training/test partitions, we extract three groups of glycans based on their ranks: high accuracy (rank 1), medium accuracy (ranks 3 and 4), and low accuracy (ranks *>* 6). (a) Representative motifs enriched for each group, based on lowest binomial test p-value. No statistically significant motifs were found for the low accuracy O-glycans due to small sample size. (b) Random sample glycans from each group. (c) Overlapping histograms of prediction ranks for tested N and O glycans (blue and yellow, respectively, with brown representing overlapping bars of the histograms). A random sample of correct and predicted glycan structures may be found in Figure S-2.

### The model generalizes to CHO cell glycans

The primary value of a machine learning model is its ability to generalize to new contexts. We have thus far only demonstrated our model’s accuracy on CFG array glycans. These arrays house a diverse set of glycans that could appear in many different organisms and contexts.

By contrast, glycans restricted to a single species or cell type will display many constraints on their structure,^39–41^ and may carry motifs underrepresented in the CFG array, which can affect the accuracy of our model. Accordingly, we sought to test our model on glycans from both recombinant and naturally-occurring glycoproteins produced in CHO cells.

From a pharmaceutical point of view, this question matters because CHO cells are the most commonly used cells in protein drug production.^36^ A carefully curated list has been reported, which contains all N-linked glycans that have thus far been reported on proteins produced by CHO cells in meaningful quantities.^37^ Will a Boltzmann model trained on CFG glycans be able to sequence these CHO cell glycans?

Since we did not have lectin binding data on all of the CHOGlycoNET glycans, we could not straightforwardly answer this question. However, we found that 25 CHOGlycoNET glycans overlapped with the CFG dataset, and we thus had lectin profiles for them. We therefore took these to be our test set, and took all the CFG glycans excluding these to be our training set (Figure 6). The model does not see any of these CHO glycans in its training inputs.

**Figure 6:**
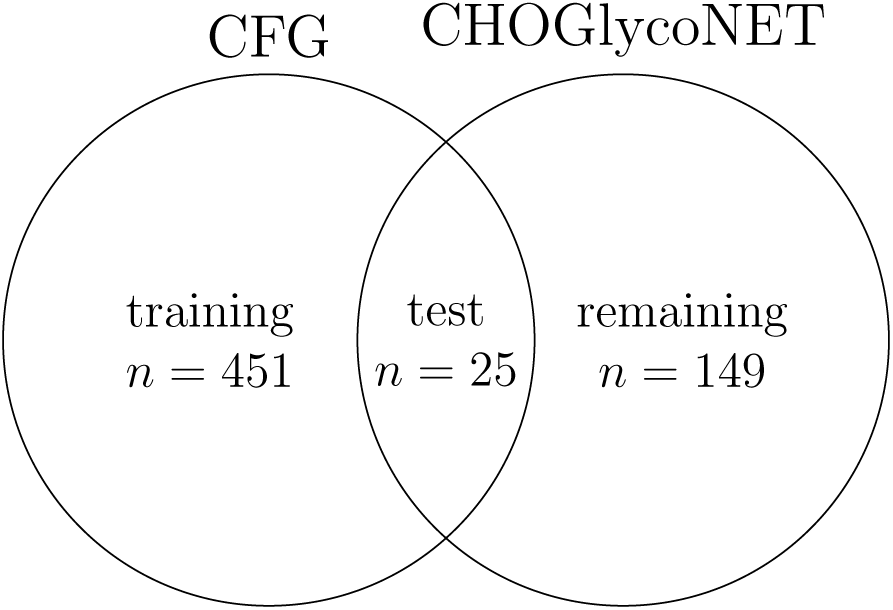
Training and test partitions of CFG and CHOGlycoNET glycans. *n* denotes the number of glycans in each set.

Though these 25 glycans do not constitute a random sample of CHOGlycoNET glycans, when comparing these glycans to other CHOGlycoNET glycans, we did not observe any major differences, and we believe this sample to be reasonably representative of the whole. Indeed, the CHOGlycoNET set is covered by the Markov model generated from our CFG glycans, which is further evidence that the untested CHOGlycoNET glycans are not very different and would be equally well predicted if we had the lectin binding data on them.

Our model was remarkably predictive of the CHOGlycoNET glycans, exactly identifying 14 / 25 of glycans, and placing 23 / 25 glycans in the top 3 predictions (Figure 7). This could be due in part to the reduced complexity of N-glycosylation when confined to a single cell type. If the model were also trained to take context or species of origin as input, we would expect to witness even greater accuracy.

**Figure 7:**
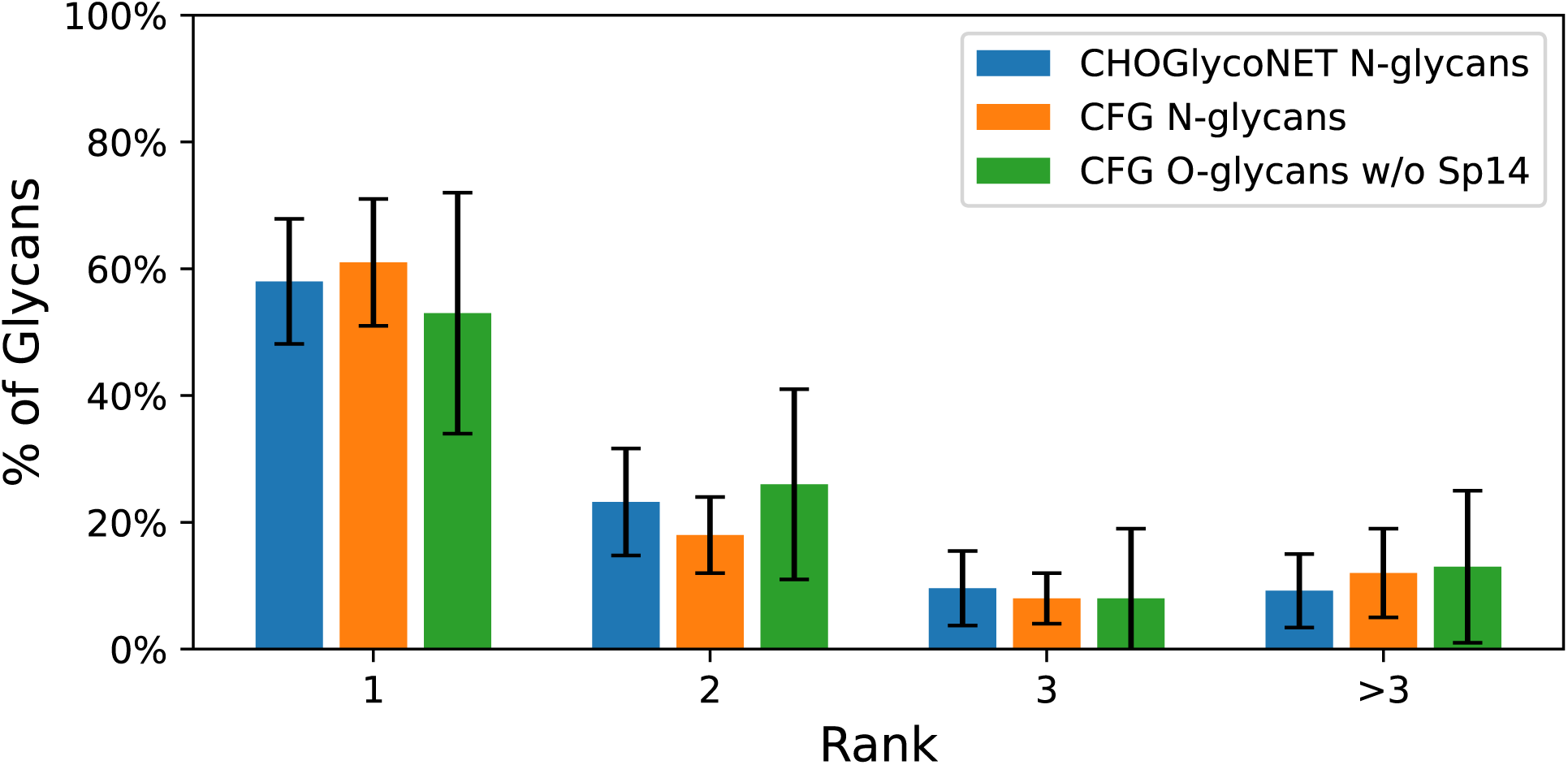
Ranks returned for each correct CHO N-glycan alongside the accuracy for general CFG N- and O-glycans. The CFG data are taken from the experiments of the previous section, where an 80/20 train/test partition was used. Bernoulli *σ* is used for the CHO error bars, and standard deviation is used for the CFG error bars. Note that the CHO results closely match the CFG N-glycan results, implying that the model generalizes well.

### Some motifs are invisible to lectin binding

Not all common glycan motifs can be identified with the set of lectins we studied here. In general, terminal motifs were the most easily identified, presumably because they are the least susceptible to steric interference. This is in broad agreement with most known lectin binding rules.^24,26,42,43^

We observed, however, that there are also terminal motifs that were less well-identified, and finding GBPs that recognize these features could improve performance of our model in future studies. The following is a list of motifs that we believe may benefit from identification of new partner GBPs:

- Terminal Neu*α*3Gal, for which GBPs have been reported but were absent in our dataset.^43–48^ By contrast, Neu*α*6Gal, is one of the most sharply identified motifs in our dataset (Figure S-3), demonstrating one of the key advantages of GBPs: *bond specificity*.
- Terminal Fuc*α*2Gal*β*, which is recognized in some contexts (i.e. BgH) but not in others.
- Internal polyLacNAc, which is recognized by several GBPs, but not reliably.

As noted, some motifs can already be recognized by commercially-available lectins absent in our data. Others could potentially be identified using engineered glycan-binding molecules such as nanobodies.^49–52^ By progressively illuminating these dark corners of the motif space we can further enhance the information content of our array.

### Robust motif binding is rare among lectins

The binding motifs of many lectins have been studied and catalogued, ^24–27,43^ and experiments have been performed to evaluate the relative affinities of lectins for some glycoproteins of interest.^53–55^ However, little has been published to quantify the specificity of lectin binding, and such studies have only been done on a predetermined set of known glycan motifs.^42,56^ Here, we extract all statistically significant motifs (see Methods - Motif recognition), and quantify the specificity of lectin binding for each of these motifs using the mutual information, which has several desirable properties (see Methods - Mutual information).

In our analysis, we find a wide range of lectin accuracies, with only a handful of lectins binding any motif robustly (Figures 8, S-3, S-7), consistent with an earlier study.^56^ Figure 8 shows the best lectins for each significant motif in the cases of N- and O-glycosylation.

**Figure 8:**
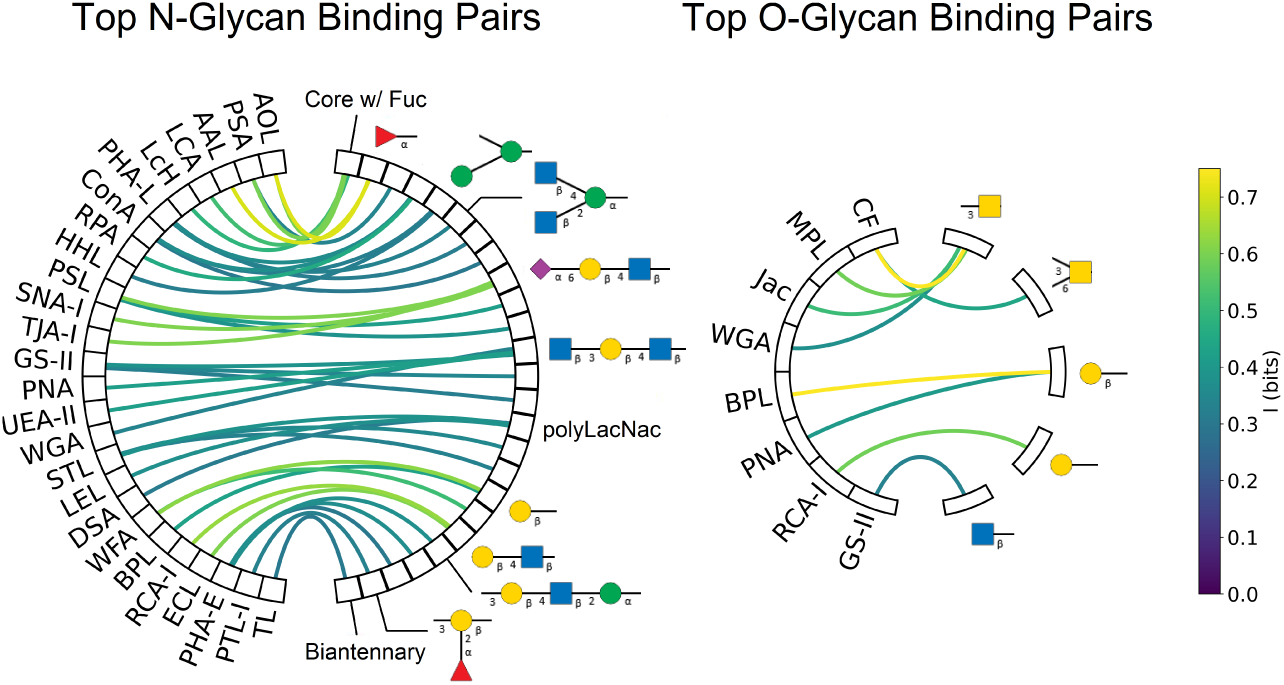
Mutual information between lectins and motifs. Only pairs with *≥.*3 bits of information are displayed. Each box on the right corresponds to a motif, but to save space only the highest correlation motif is displayed for each lectin.

### Most lectins only reliably bind one motif

Lectins often have secondary and tertiary binding motifs, i.e. additional motifs that lectins may recognize with less affinity or frequency than their primary binding motifs.^24,42^ To test if these secondary motifs are predictive of lectin binding, we measured the increase in mutual information gained by adding additional motifs. On average, we found that over 80% of the information captured by a lectin comes purely from its primary binding motif, with additional motifs contributing little extra information (Figure 9).

**Figure 9:**
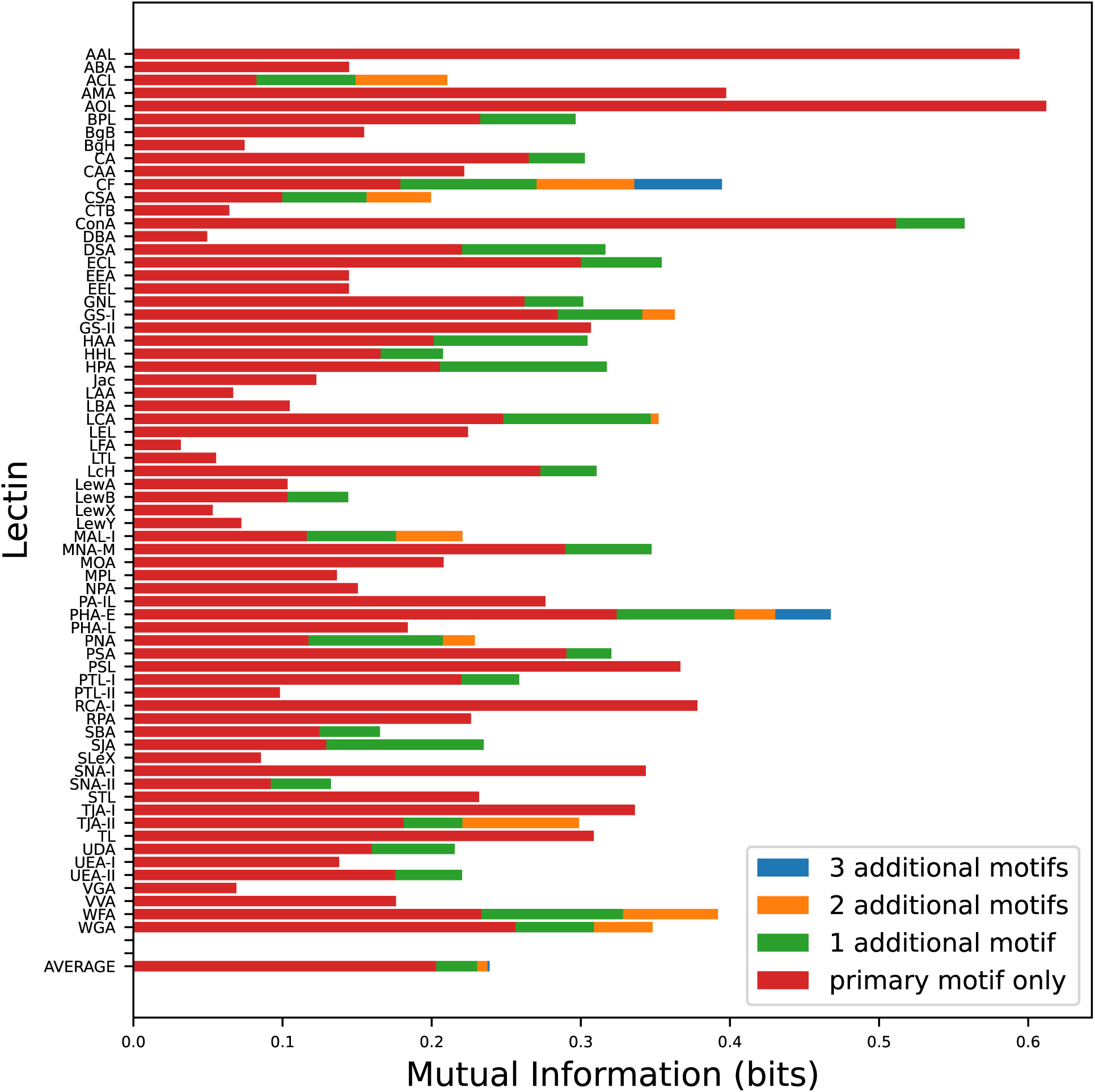
Mutual information between lectins and their best-binding motifs. On average, 82% of a lectin’s captured information comes from its primary binding motif. This increases to 86% and 89% when restricting to N and O glycans respectively, due to increased specificity.

This fact has positive and negative implications with regard to using lectins to determine glycan structure. On the one hand, it is unlikely that a single lectin can be used to gather information on the presence of multiple motifs. On the other hand, it makes endeavors to interpret lectin binding results much simpler, particularly for a machine learning algorithm. Our results are consistent with the conclusion that any secondary binding motifs that exist are too closely correlated with their primary binding motifs to generate much extra predictive power. It is also possible that there are secondary motifs that do not appear in sufficient quantity in our dataset to be statistically significant, but would become relevant in datasets where they are more common. In either case, for most lectins, primary binding will capture most of the available information.

## Discussion

In this study we analyzed the information content of lectin binding, and demonstrated its utility for *de novo* glycan sequencing. We have shown that the sequencing of CHO N-glycans should be possible in theory with current lectin panels, and that a broader range of glycans can likely be sequenced if care is taken to curate a diverse set of motif-specific GBPs. We have also shed light on which lectins are most dependable, and which motifs are least identifiable, both of which could aid future researchers interested in using lectins for their studies. Our analysis is very general and can be used as a reference for choosing the best set of lectins for any task.

A straightforward way to improve upon our results would be to utilize a more comprehensive lectin panel. By identifying the motifs least recognized by the public lectin set studied here, we provide a roadmap for future researchers to improve upon our results. For some of the more challenging motifs we identified, GBPs may already exist. For the rest of the motifs, our analysis could guide the engineering or search for new GBPs.

With modern synthetic glycobiology techniques, methods are emerging that are increasingly capable of engineering macromolecules to bind specific glycan moieties. ^50–52,57,58^ Aptamers, lamprey antibodies, and glycan-binding nanobodies are just a few examples of molecules that can be employed to assay glycan motifs.^49,59–62^ We hope that our identification of motifs that are difficult to detect can shine a spotlight on fruitful future endeavors in this field.

Since our results were obtained by measuring lectin binding on glycan arrays, our findings should apply broadly to the gamut of lectin array techniques available today. Microfluidic integration and other refinements have made these microarrays high-throughput, reproducible, and tailorable to many lectin sets.^63–66^ An intriguing alternative approach would be to use multiplexed DNA-barcoded lectins in solution,^67^ followed by next generation sequencing to quantify lectin binding. This would have the benefit of presenting lectins in solution, which avoids the problem of array presentation seen in our O-glycan data analysis and allows for flexible use cases.

A critical next question to answer is whether this technology can be used to resolve mixtures of glycans into their component parts. While this problem may seem highly under-constrained, there are many biological constraints on glycan synthesis that narrow the set of plausible glycoprofiles.^39–41^ Often, there will only be a few glycans present in appreciable quantities in a sample, differing only by the final monosaccharides in a chain. Therefore, it is not inconceivable that lectins in a biological context could be sufficiently accurate for this task.

Despite its ubiquity in nature, glycobiology has long remained insulated from the broader scientific community. It is our hope that by making glycoprofiling more efficient and accessible to non-experts, we can better share what our field has to offer and reap the rewards of integrative science. Glycosylation should not be seen as an excess complication, it should be seen as an opportunity for discovery.

## Methods

### Data processing

For this analysis, we utilized published lectin binding data from CFG arrays.^24^ Glycans from the CFG Mammalian Glycan Array were filtered to contain only fucose (Fuc), galactose (Gal), N-acetylgalactosamine (GalNAc), glucose (Glc), N-acetylglycosamine (GlcNAc), mannose (Man), and sialic acid (Neu) residues, since these were the only monosaccharides in sufficient abundance in the CFG array for meaningful statistics to be extracted.

Additionally, we discarded glycans smaller than 2 monosaccharides, and we ignored modifications such as sulfurylation and phosphorylation. Although these modifications are known to impact lectin binding,^42,52^ they were not common enough in our data for a statistical analysis. Incorporation of these features into future studies should result in greater glycan resolution.

### Motif recognition

Motifs were taken to be any connected subgraph of any glycan which appeared more than 5 times in our dataset. These directed subgraphs are imagined to include their incoming and outgoing edges as well, so each motif includes the number of unspecified substitutions at each residue. In other words, we include and differentiate between both terminal motifs like Gal*β*1-4GlcNAc*β*-X and internal motifs like X-Gal*β*1-4GlcNAc*β*-X. Motifs of size *>* 5 monosaccharides were removed to reduce computational load. Inclusion or exclusion of these large motifs was not found to impact performance.

### Binarization of lectin binding

Since lectin binding strength measured in RFU can vary greatly depending on context for a given motif, linear binding measures are not appropriate (Figure S-4). Instead, we binarize our lectin binding data by generating a logarithmically-spaced range of thresholds from 1000 to 20,000 RFU to serve as binding cutoffs. For each cutoff, if the raw binding value exceeds the threshold, we record this as a discrete binding event. Each lectin-glycan pair will therefore have several associated 1s and 0s to denote the thresholds surpassed by the binding.

### Markov modelling of glycan space

Markov models of varying complexity have been used previously in glycobiology.^41,68–70^ Here we use a simple model that takes the previous two residues in a chain and predicts the next one. This is less sharp than other models which have been used because the goal is not to exactly model the glycan space, which may be constrained in many ways in a given species, but to provide a very diverse range of possible glycans. This will allow us to measure the *de novo* sequencing accuracy, and later constrain the space further in a cell or species specific manner to obtain greater accuracy.

We found our model to produce reasonable and diverse results (Figures S-5 and S-6), and calculated its entropy to be 12.3 bits based on the CFG glycan set. This implies that a random sample of on the order of 10^4^ glycans should provide good coverage of the full range of possible CFG-like glycans. An even more general model would take only a single monosaccharide as input, but we found this to produce pathological results (Figure S-5).

### Model training

Since our Boltzmann model is convex, all training procedures should converge to the same unique maximum likelihood solution. We used gradient descent with step size 0.1. We did not find L1 or L2 regularization terms to make a meaningful difference in our results, so we did not utilize them in any of the figures presented here.

### Mutual information

One way to measure the sharpness of a lectin at recognizing a motif would be to use the correlation coefficient; however this suffers from three drawbacks: First, a linear measure is not appropriate because the binding strength can vary wildly for the same motif in different contexts. Second, a high correlation coefficient can be achieved when a rarely binding lectin overlaps by chance with a rarely expressed motif. It would then require further parameters (p or *σ* for example) to determine whether this binding is actually statistically significant. Third, in a scenario we examine in our paper, if a lectin binds multiple motifs it may not be well correlated to any one of them.

One measure that overcomes these deficiencies is the mutual information:

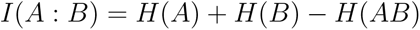

where *H* is the Shannon entropy.

This measure has many desirable properties: First, it is measured in bits, enabling us to compare the amount of information obtained to the amount of information left unknown. Second, *H*(*A*) *≥ I*(*A* : *B*) *≤ H*(*B*). This means that rare, statistically insignificant motifs yield insignificant mutual informata. Third, A and B need not be a single lectin or motif. This will allow us to assess the impact of binding patterns involving more than one lectin or motif.

### Cross entropy

Any probability model’s accuracy can be assessed using the cross entropy. If we restrict our attention to the problem of predicting the presence or absence of a single glycan motif, then our classification is binary and the cross entropy loss reduces to the logistic loss function:

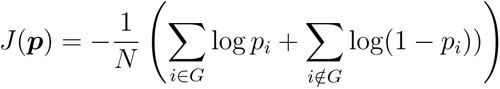

where *p_i_*denotes the model’s predicted probability that glycan *i* possesses the desired motif, *G* denotes the set of glycans with said motif, and *N* denotes the total number of glycans.

We consider the case of using any two lectins to predict a given motif. We may define zeroth, first, and second order models as follows:

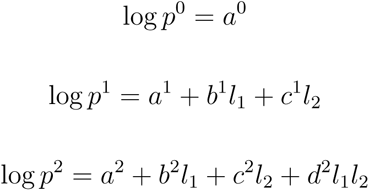

where *l*_1_ and *l*_2_ denote the lectin binding binaries and we optimize the parameters *a, b, c, d* to minimize the logistic loss.

Our models are obtained by optimizing these parameters. The zeroth order model takes no lectin inputs, so its loss is simply the motif’s entropy *H*. We may quantify the amount of information a model can extract by improvements Δ*H* to this loss. The differences Δ*H*_1_ = *H − J* (*p*^1^) and Δ*H*_2_ = *H − J* (*p*^2^) correspond to the amount of information extracted about the motif by the first and second order models, respectively. As we show in Figure 2, Δ*H*_2_ *−* Δ*H*_1_ is quite small, so the models capture a similar amount of information from the lectin signal.

## Supporting information available

The following supporting files are available free of charge:

- Supplemental Table n Figures.pdf - This file contains a lectin code glossary, samples from our model, and some additional plots of information parameters we analyzed.
- pickle files.zip - This file contains the raw lectin binding data used for our analysis, and other relevant information.

## Acknowledgements

This work was supported by funding from the Novo Nordisk Foundation (NNF20SA0066621), the National Institute of General Medical Sciences (R35 GM119850), and support from a UCSD Academic Senate Award (RG102546).

## Conflict of interest disclosure

N.E.L. and A.W.T.C. have submitted patents associated with the use of lectin binding patterns for determining glycan structures and disease diagnostics. N.E.L. is also co-founder and holds financial interest in NeuImmune and Augment Biologics, which focus on glycoprotein therapeutics.

## TOC Graphic

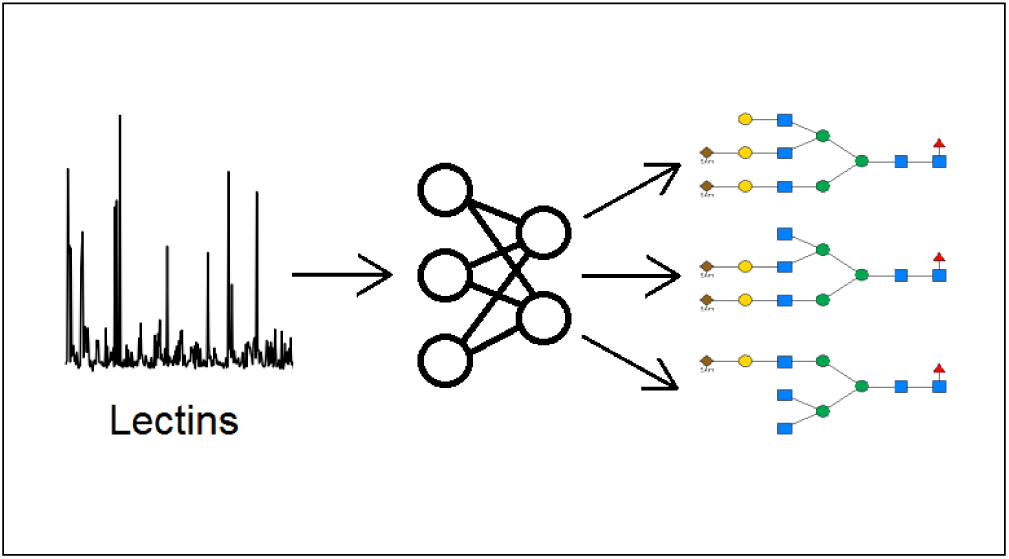

## Supporting information

**Table S1:**
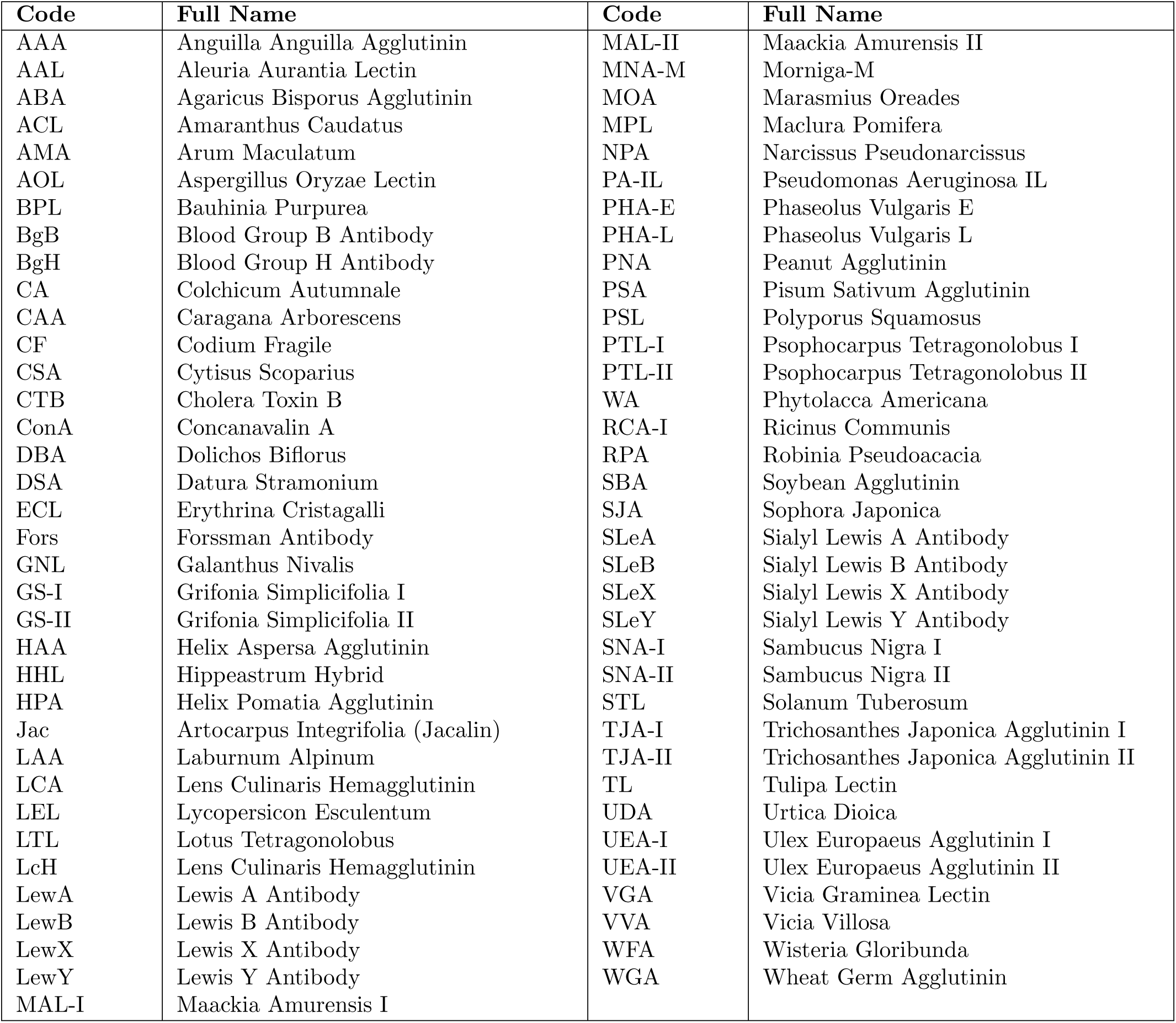
Lectin code glossary.

**Figure S-2:**
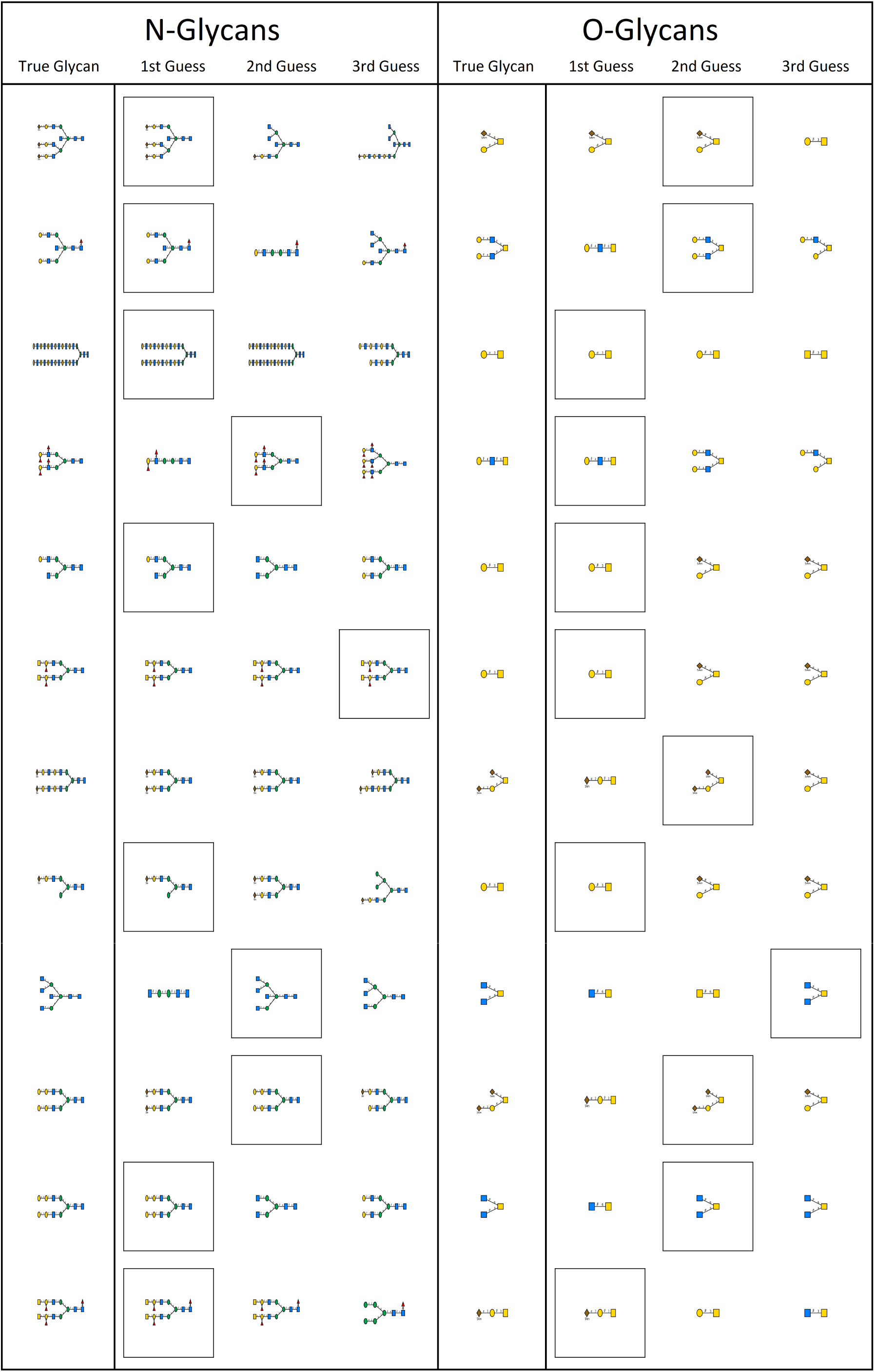
Random sample of N and O glycans, along with our model’s predictions based on their respective lectin profiles. The correct glycan is boxed. Note that even when the correct glycan is not the model’s first pick, it is often in the top three predictions. Also, the predicted glycans a^S^r^2^e typically very similar to the correct glycan, particularly in terms of their terminal motifs.

**Figure S-3:**
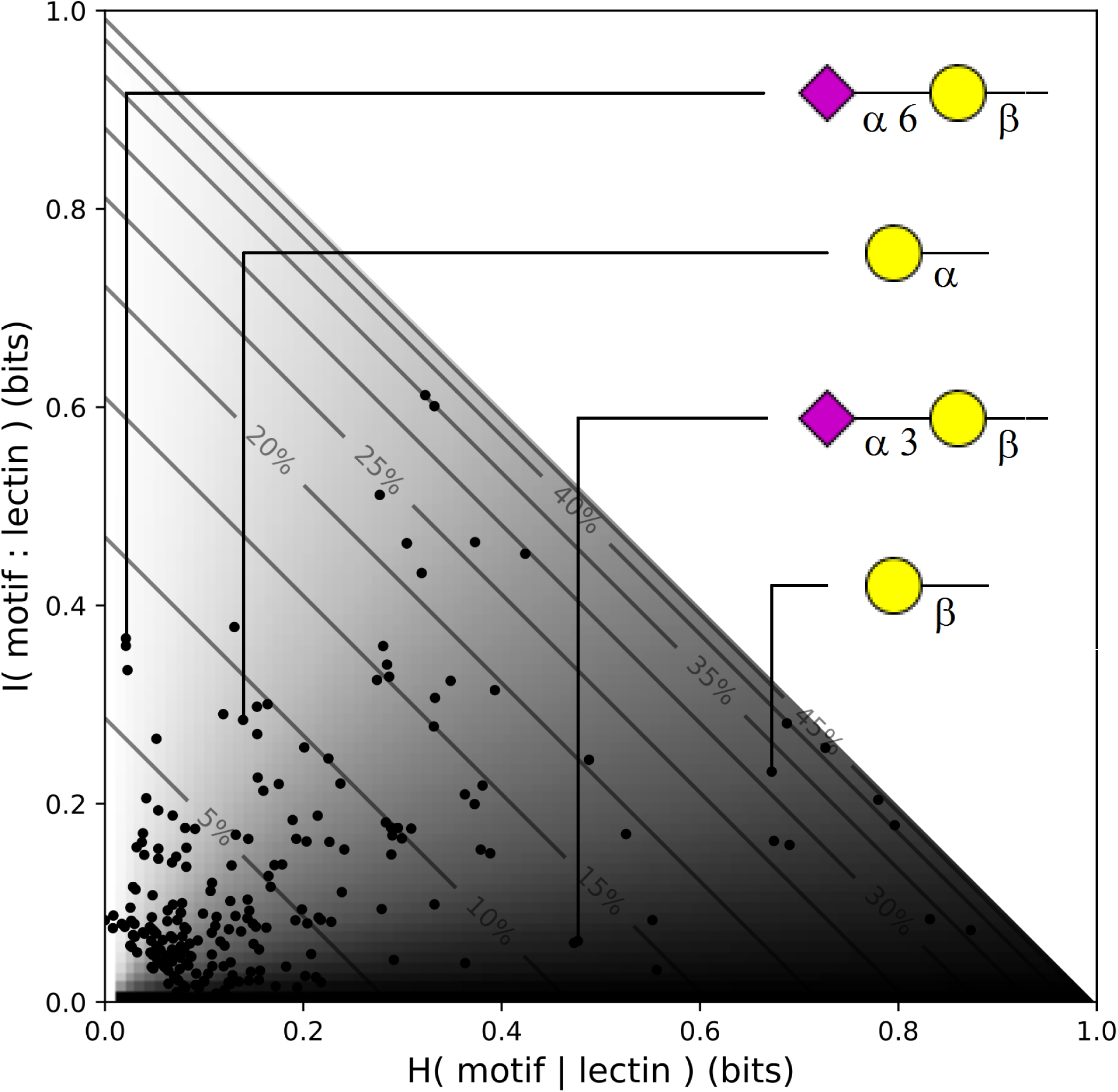
Scatter plot of mutual information and conditional entropy for each motif and its best binding lectin. When mutual information is high, the lectin is a strong predictor of the motif’s presence. When conditional entropy is high, there remains a great deal of uncertainty in whether or not the motif is present, even after the information from lectin binding is taken into account. Note that the sum *I*(*m* : *l*) + *H*(*m|l*) = *H*(*m*) is the motif entropy, so the ratio between *I*(*m* : *l*) and *H*(*m|l*) is a measure of the sharpness of the binding. Motifs on the light side are well-recognized, while motifs on the dark side are invisible to the set of lectins studied here. Contours correspond to the prevalence of each motif in our dataset, with the least common motifs occurring in the bottom left corner. Note that variations in bond orientation can have dramatic impacts on motif recognition.

**Figure S-4:**
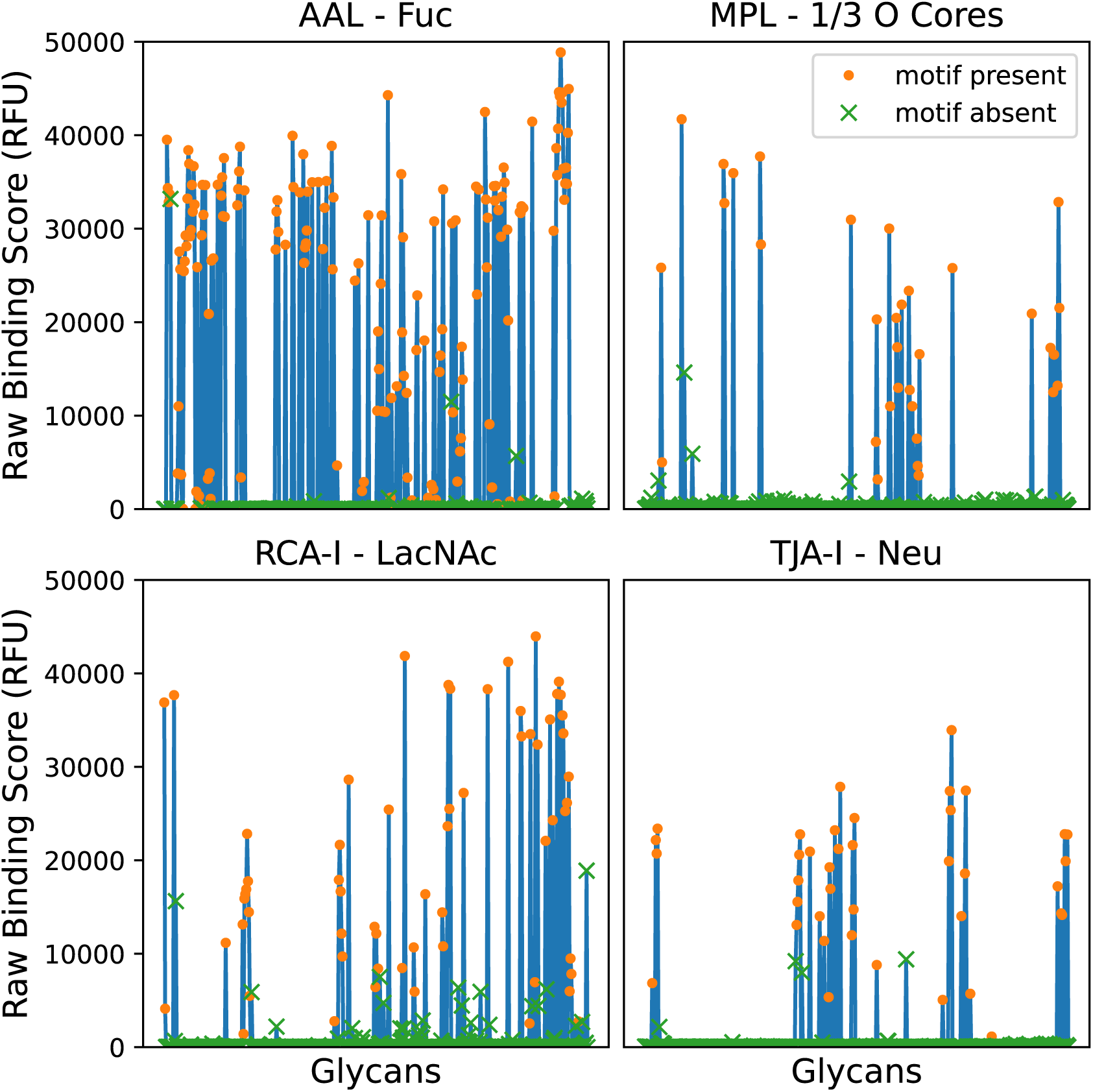
Raw binding values from our CFG arrays for four of our sharpest lectin-motif pairs. AAL binds Fuc*α*, MPA binds O glycan cores 1 and 3. RCA-I binds terminal Gal*β*4GlcNac, and TJA-I binds Neu*α*6Gal*β*4GlcNAc. Note that even among cases when the motif is present, binding strength can vary by a factor of 2 of more. Also note that the apparent lack of variance in AAL binding strength is due to saturation.

**Figure S-5:**
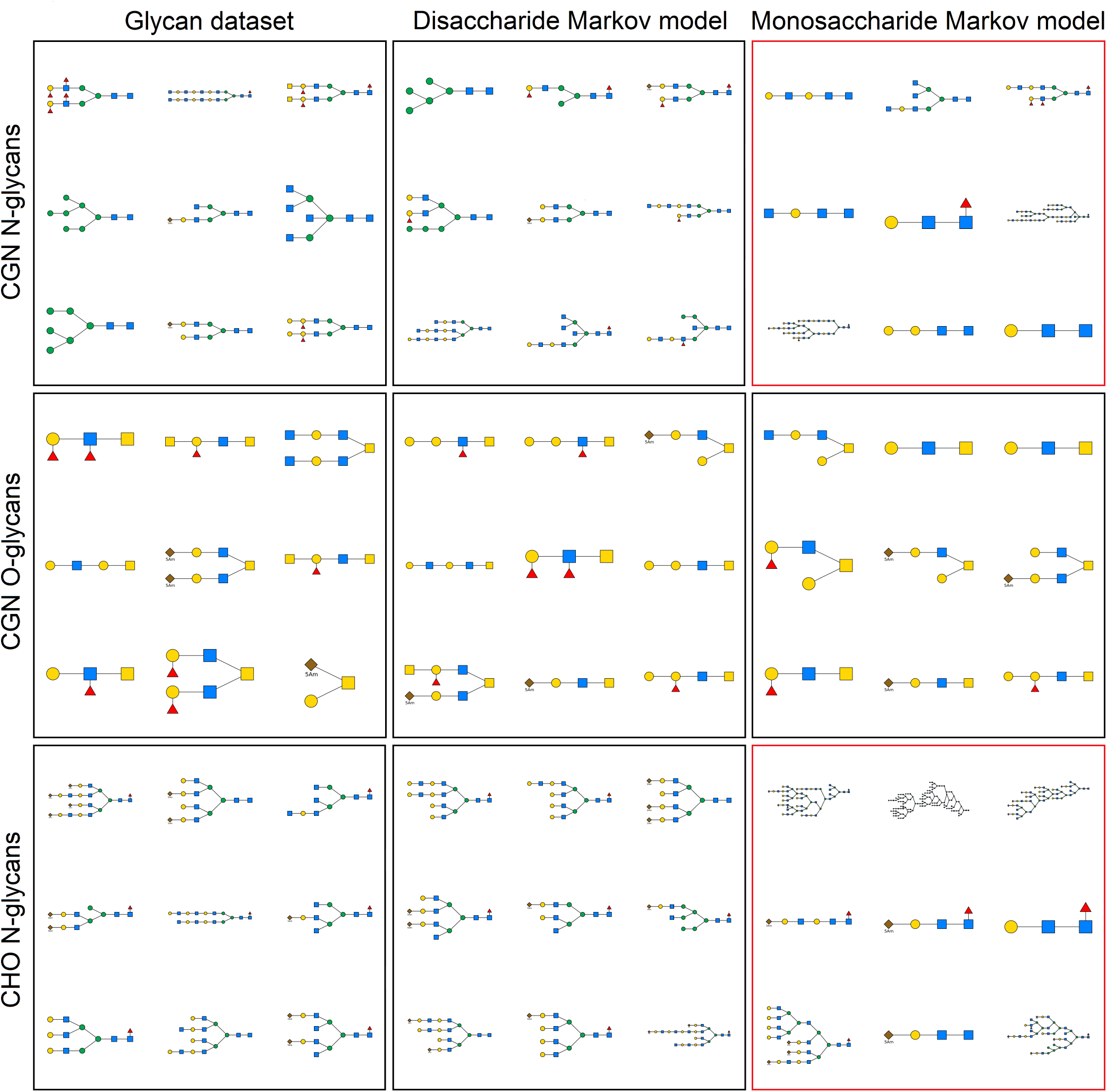
Random sample glycans from each of our datasets alongside random samples from their respective Markov models. Note that while the disaccharide model used in our paper appears to model the glycans well and produce diverse results, the more general monosaccharide model produces pathological samples on both N-glycan sets.

**Figure S-6:**
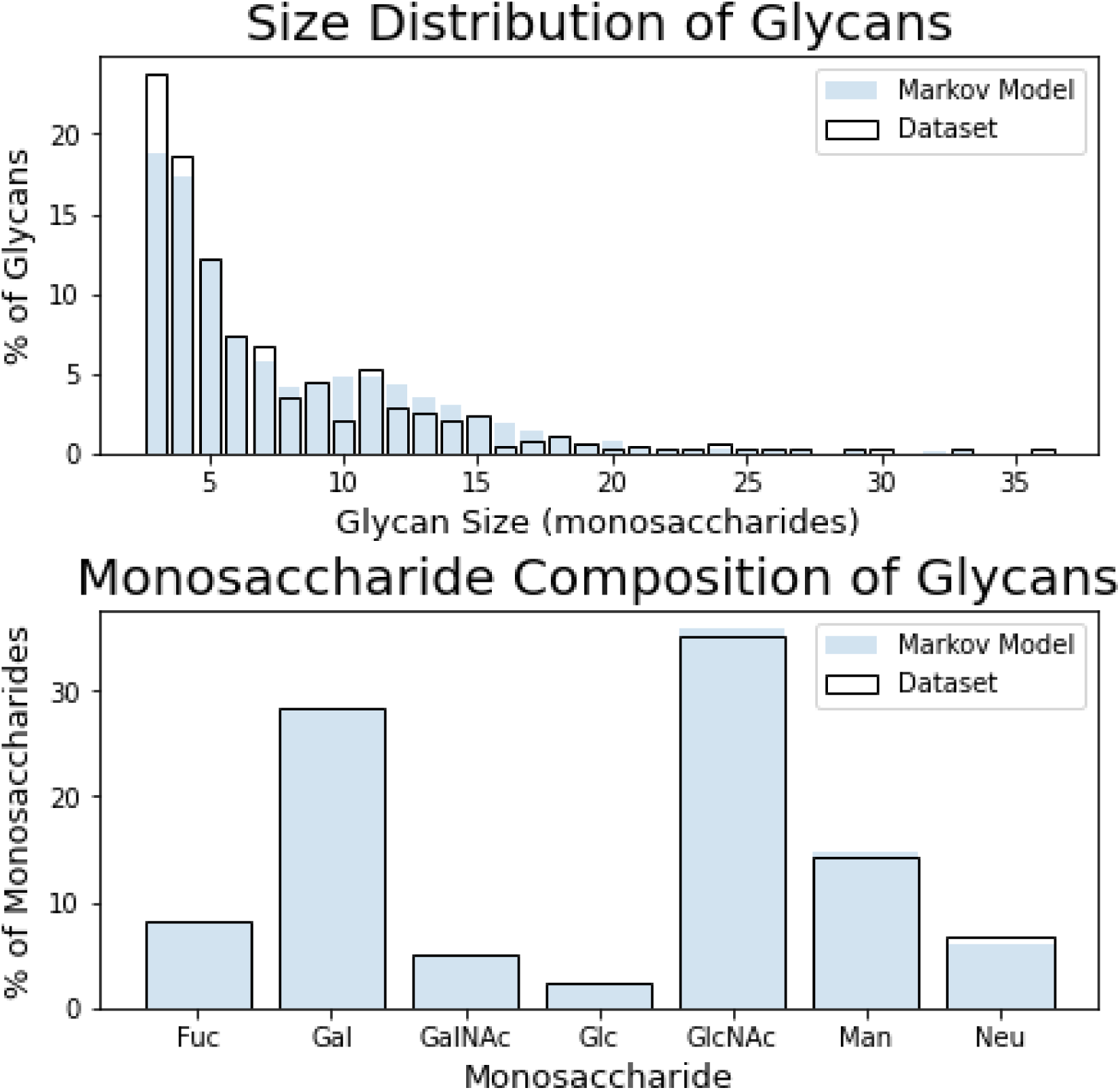
10,000 samples from Markov model of CFG glycan set. Our model appears to generate glycans of the right size and monosaccharide composition.

**Figure S-7:**
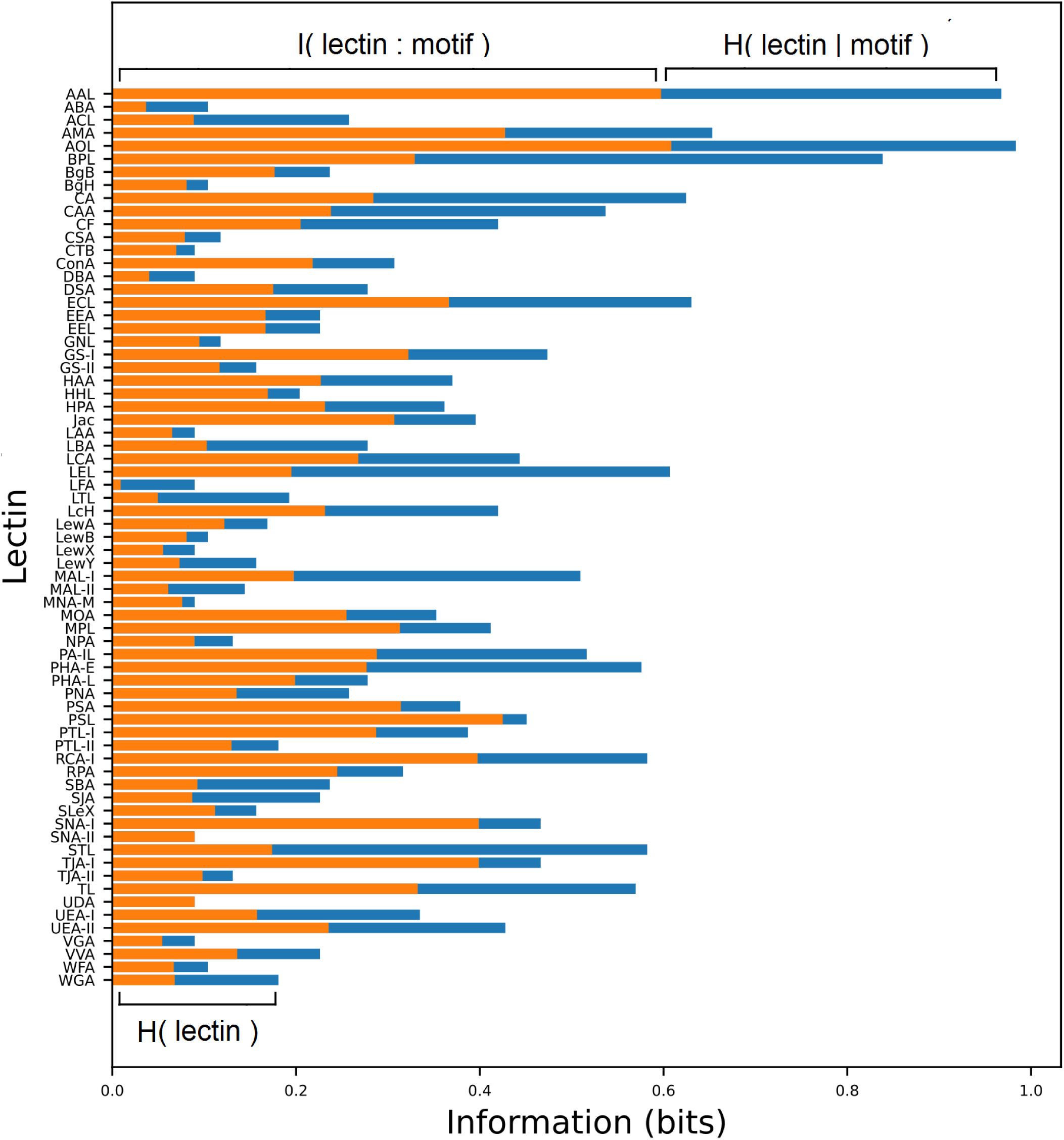
Information extracted by each lectin about its most sharply bound motif. When the mutual information is much larger than the conditional entropy, the binding is very specific. Note that in most cases, either the conditional entropy is high, implying that the binding is not very robust, or the total entropy is low, implying that the relevant motif is not very common.

## Notes

### Competing Interest Statement

The authors have declared no competing interest.

### Summary of Updates

By removing the mispresented Sp14 spacer, we were able to improve accuracy on O-glycans. We also made several other changes to enhance the usefulness and readability of the manuscript.

