## Supplementary material for "A Boltzmann model predicts glycan structures from lectin binding": Data: README.txt

pickle_files.p contains a tuple of the following five variables:


(bojar_filenames, bojar_avgs, bojar_stds, rawcfgs, lectin_codes)


Here is a description of each variable:


bojar_filenames = list of filenames of raw data sets

bojar_avgs = N_lectins x N_glycans numpy array. Records mean RF value for each lectin-glycan binding pair

bojar_stds = N_lectins x N_glycans numpy array. Records uncertainty in RF value for each lectin-glycan binding pair

rawcfgs = list of N_glycans IUPAC glycan codes

lectin_codes = list of N_lectins lectin codes
